# SGK regulates pH increase and cyclin B-Cdk1 activation to resume meiosis in starfish ovarian oocytes

**DOI:** 10.1101/499699

**Authors:** Enako Hosoda, Daisaku Hiraoka, Noritaka Hirohashi, Saki Omi, Takeo Kishimoto, Kazuyoshi Chiba

## Abstract

Tight regulation of intracellular pH (pH_i_) is essential for biological processes. Fully-grown oocytes, having a large nucleus called the germinal vesicle, arrest at meiotic prophase-I. Upon hormonal stimulus, oocytes resume meiosis to acquire fertilizability. At this time, pH_i_ increases through Na^+^/H^+^ exchanger activity. However, regulation and function of this change remains obscure. Here we show that in starfish oocytes, serum- and glucocorticoid-regulated kinase (SGK) is activated by the PI3K/TORC2/PDK1 signaling after hormonal stimulus, and is required for the pH_i_ increase and cyclin B–Cdk1 activation. Furthermore, when we clamped pH_i_ at 6.7, corresponding to the pH_i_ of unstimulated ovarian oocytes, hormonal stimulus normally induced cyclin B–Cdk1 activation; thereafter, oocytes initiated germinal vesicle breakdown (GVBD), but failed to complete it. Thus, SGK-dependent pH_i_ increase is likely prerequisite for completion of GVBD in ovarian oocytes. We propose a model that SGK drives meiotic resumption through concomitant regulation of pH_i_ and the cell-cycle machinery.

## Introduction

Intracellular pH (pH_i_) is tightly regulated in living cells. Increases in pH_i_ are required for a variety of physiological and pathological processes, including early embryonic development^1,2,3^ and cancer cell survival^4^. A key pH_i_ regulator, the sodium–proton exchanger (NHE) is a 12-transmembrane protein that increases pH_i_ by exporting intracellular H^+^ and importing extracellular Na^+^ (ref. ^5^). Many kinases, including serum- and glucocorticoid-regulated kinase (SGK)^6-8^, p90RSK^9^, and p38MAPK^10,11^, activate NHE in mammalian cells.

An NHE-dependent pH_i_ increase has also been reported during oocyte maturation^12-17^, a fundamental process involved in the generation of fertilizable eggs, although the underlying regulatory mechanism and function of pH_i_ increase during oocyte maturation are not completely understood. In the ovaries of most animals, fully-grown oocytes, which have a large nucleus called the germinal vesicle (GV), are arrested at prophase of meiosis I (ProI)^18^. Upon extracellular stimuli by maturation-inducing hormones, ovarian oocytes resume the meiotic cell cycle via activation of cyclin B–Cdk1 and subsequently undergo germinal vesicle breakdown (GVBD)^19-21^. Subsequently, oocytes undergo another arrest at the metaphase of meiosis I or II (MI or MII arrest) that enables successful fertilization^22^. Starfish oocytes have been used as a model to study oocyte maturation for more than half a century^23^. Previously, we reported that pH_i_ of ProI-arrested ovarian oocytes is low (~6.7) due to the relatively high CO_2_ and low O_2_ concentrations in the body cavity of female starfish^17^. Soon after stimulation by the hormone 1-methyladenine (1-MA)^24^, pH_i_ increases to ~6.9^17^, after which the oocytes undergo GVBD and ultimately arrest at MI^16,17^. To maintain the MI arrest, pH_i_ is maintained at ~6.9 until spawning^16,17,25-28^. The increase in pH_i_ following 1-MA stimulus is known to require starfish NHE3 (sfNHE3)^16,17,25^. However, the upstream signaling and function of this pH_i_ increase remain elusive.

For analysis of pH_i_ regulation, unstimulated starfish oocytes are isolated from ovaries, placed in artificial seawater (ASW), and used for experiments because various techniques, such as microinjection, are easily applicable to the isolated oocytes. ASW contains relatively low CO_2_ and high O_2_ relative to the inner body cavity^17^. Due to this difference in gas concentrations, the basal pH_i_ of isolated oocytes is ~7.0, ~0.3 units higher than that in ovarian oocytes^17^. Soon after 1-MA treatment, this value increases further, to ~7.3, in a sfNHE3-dependent manner^16,17,25^. 1-MA induces Gbg-dependent activation of phosphoinositide 3-kinase (PI3K)^29-33^. Previously, we demonstrated that the pHi increase is mediated by Gbg and PI3K^16^, although the molecular link between PI3K and the sfNHE3-dependent pH_i_ increase remains unclear.

One candidate for this link is SGK, which is activated downstream of PI3K and upregulates NHE3 in mammalian cells^6,34-36^. Mammalian SGK, which belongs to the AGC kinase family^37^, has three isoforms: SGK1, 2, and 3^35,38,39^. All three proteins have an activation-loop (A-loop) and a C-terminal hydrophobic motif (HM)^35^, but only SGK3 has the phospholipid-binding domain called the Phox homology domain (PX domain)^39^. Upon agonist stimulation, the HM is first phosphorylated by mammalian target of rapamycin complex 2 (mTORC2) in a PI3K-dependent manner^38-40^. Phosphoinositide-dependent kinase 1 (PDK1) interacts with the phosphorylated HM and phosphorylates the A-loop^41^, resulting in full activation of SGK^35,36,42,43^. Although it is not known whether SGK is activated in starfish oocytes, we previously showed that the C-terminus of sfNHE3 has a putative consensus sequence for phosphorylation by SGK, and can be phosphorylated by recombinant human SGK1 *in vitro*^25^. Furthermore, PDK1 and TORC2 are functional in starfish oocytes, in which they activate another AGC family kinase, Akt^37^, through phosphorylation of its A-loop and the HM^44,45^. In this context, Akt participates in inhibition of Myt1 and activation of Cdc25^32,45-47^, leading to cyclin B–Cdk1 activation through dephosphorylation of Cdk1 at Thr14 and Tyr15^23,47^. Collectively, these observations inspired us to investigate whether SGK serves as a downstream mediator of PI3K for the pH_i_ increase in starfish oocytes.

Another issue that remains to be resolved is the role of the pH_i_ increase following 1-MA stimulation. Previously, we established a method for clamping pH_i_ at a desired value in starfish oocytes^17^, enabling us to examine the effects of altered pH_i_. In this study, to elucidate the regulation and function of the pH_i_ increase in oocytes, we examined the involvement of SGK in the pH_i_ increase following 1-MA stimulus and the effects of pH_i_ alteration on meiotic resumption. We found that SGK activation is prerequisite for two mutually independent events: pH_i_ increase and cyclin B-Cdk1 activation. Moreover, by clamping pH_i_ at various values, we showed that oocytes exhibit defective GVBD at reduced pH_i_ values. Based on these findings, we propose a model for SGK-dependent meiotic resumption in starfish ovarian oocytes.

## Results

### Starfish SGK is a homolog of human SGK3

To investigate the possible involvement of SGK in the pH_i_ increase of oocytes during hormonal stimulation with 1-MA in the starfish *A. pectinifera*, we cloned the cDNA of starfish SGK (sfSGK). The open reading frame (ORF) of the cDNA encodes a polypeptide of 489 amino acids with a predicted molecular mass of 56 kDa. At the amino acid sequence level, sfSGK is 52% identical to the human SGK3 PX domain (Fig. 1a, blue open box) and 77% identical to the human SGK3 catalytic domain (Fig. 1a, orange open box). Two phosphorylation sites, which are required for activation of mammalian SGK, are conserved in sfSGK: Thr312 in the A-loop (corresponding to human SGK3 Thr320 which is phosphorylated by PDK1), and Thr479 in the HM (human SGK3 Ser486 which is phosphorylated by mTORC2) (Fig. 1a). The transcriptome database contains no other SGK isoforms for *A. pectinifera*, suggesting that SGK3 is the only member of this protein family in starfish.

**Figure 1.**
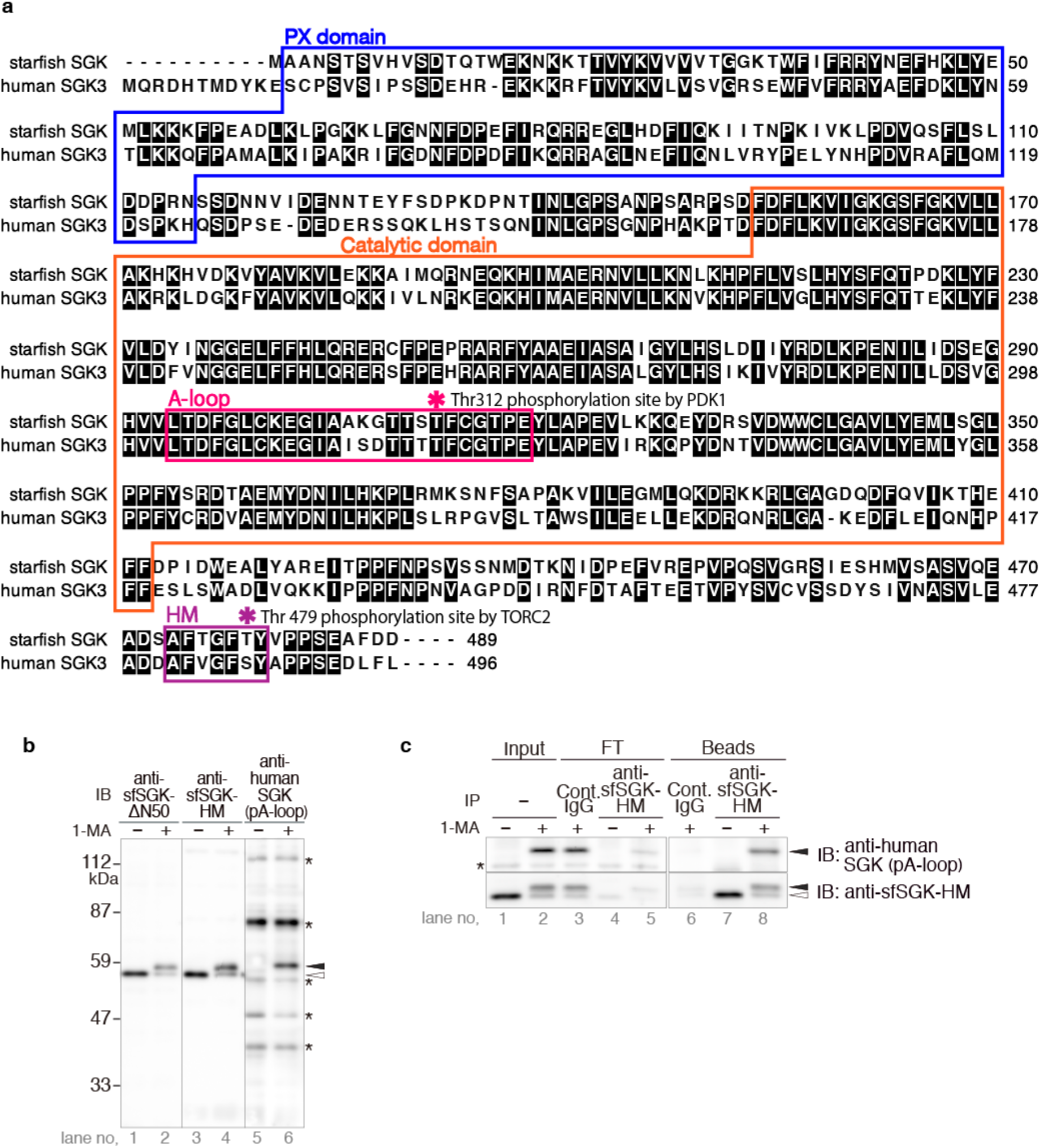
sfSGK protein and phosphorylation of its A-loop are detectable in starfish oocytes. **a** The deduced amino acid sequence of starfish SGK (sfSGK) was aligned with that of human SGK3 (NCBI accession number NP_001028750.1) using the ALAdeGAP alignment software^67^. The sfSGK sequence has been submitted to the DDBJ/EMBL/Gen Bank databases under accession number LC430700. Identical amino acids are shaded. Dashes indicate gaps introduced for optimal alignment. Colored boxes indicate conserved domains: PX domain (blue), Activation-loop (A-loop; magenta), Hydrophobic motif (HM; purple), and Catalytic domain (orange). Magenta and purple asterisk indicate conserved residues phosphorylated by PDK1 (Thr312 in starfish; Thr320 in human) and TORC2 (Thr479 in starfish: Ser486 in human), respectively. **b** Unstimulated starfish oocytes were incubated with or without 1-MA for 4 min, followed by immunoblotting with anti-sfSGK-N50, anti-sfSGK-HM, and anti-human phospho-SGK (anti-human SGK [pA-loop]) antibodies. Asterisks indicate non-specific bands. The result shown is representative of three independent experiments. **c** Immunoprecipitation (IP) was performed with anti-sfSGK-HM antibody or control IgG using extracts prepared from unstimulated or 1-MA stimulated oocytes. Input extract (Input), flow-through (FT), and beads (Beads) samples were analyzed by immunoblotting with the indicated antibodies (IB). Asterisk indicates non-specific bands detected by anti-human phospho-SGK (anti-human SGK [pA-loop]). The indicated result is representative of two independent experiments. Closed and open arrowheads indicate positions of the upper and lower bands of sfSGK, respectively (**b**, **c**).

For this study, we generated two types of antibodies: one against a recombinant fragment of sfSGK lacking the N-terminal 50 amino acids (anti-sfSGK-ΔN50), and the other against a C-terminal peptide (17 amino acids) containing the HM of sfSGK (anti-sfSGK-HM). We evaluated the reactivity of these antibodies by immunoblotting starfish oocytes. Both antibodies detected a protein of 56 kDa, corresponding the predicted molecular mass of sfSGK in unstimulated oocytes (Fig. 1b, *left* and *middle* panels), suggesting that these antibodies recognized a sfSGK protein. The sfSGK band underwent a mobility shift after 1-MA treatment (Fig. 1b, *left* and *middle* panels), suggesting that the protein is phosphorylated after 1-MA stimulation.

Full activation of mammalian SGK is achieved by PDK1-dependent phosphorylation of A-loop^35,41^. The amino acid sequence of the A-loop in mammalian SGK is quite similar to that of sfSGK (Fig. 1a). Hence, to determine whether sfSGK is phosphorylated in the A-loop in stimulated starfish oocytes, we used a commercial polyclonal anti-human phospho-SGK antibody that recognizes a phosphorylated amino acid in the A-loop of mammalian SGK. The antibody detected a ~59 kDa protein in stimulated oocytes, but not in unstimulated oocytes (Fig. 1b, *right* panel). In addition, this band ran at the same position as the shifted bands recognized by the anti-sfSGK-ΔN50 and anti-sfSGK-HM antibodies (Fig. 1b). To confirm that the protein detected by the anti-human phospho-SGK antibody was sfSGK, we immunoprecipitated sfSGK using the anti-sfSGK-HM antibody and immunoblotted with the anti-human phospho-SGK antibody. As expected, we detected protein in the bead fraction (Fig.1 c, *upper* panel, lane 8) but not the flow-through fraction (Fig.1 c, *upper* panel, lane 5), indicating that the anti-human phospho-SGK antibody reacted with sfSGK. Accordingly, hereafter we refer to the A-loop phosphorylation of sfSGK detected by this antibody as “sfSGK-pT312 (A-loop)” in figures. Taken together, these results suggest that sfSGK is phosphorylated at its A-loop, and thereby activated, in oocytes stimulated with 1-MA.

### sfSGK is activated soon after 1-MA stimulus

To compare the phosphorylation dynamics of sfSGK with Akt, Cdc25 and Cdk1, we treated isolated oocytes with 1-MA and analyzed them by immunoblotting. Mobility shift (Fig. 2a, sfSGK) and A-loop phosphorylation (Fig. 2a, sfSGK-pT312 [A-loop]) of sfSGK was detected 1 min after 1-MA stimulus. The timing of phosphorylation of Akt at Ser477 in HM by TORC2 (ref. ^45^) was similar to that of sfSGK (Fig. 2a, Akt-pS477 [HM]). The phosphorylation level of sfSGK increased rapidly, with a peak at 3 min after 1-MA treatment (Fig. 2a, sfSGK-pT312 [A-loop]), followed by a gradual decrease. A band shift of Cdc25 corresponding to hyperphosphorylation^32,47^ was detected after 7 min (Fig. 2a, Cdc25). Subsequently, activation of Cdk1 was detectable as dephosphorylation of Tyr15 (ref. ^47^) at 10 min (Fig. 2a, Cdk1-pY15), followed by onset of GVBD at approximately 17 min. These results indicate that sfSGK was rapidly activated at the same time as Akt, earlier than Cdc25 hyperphosphorylation and cyclin B–Cdk1 activation.

**Figure 2.**
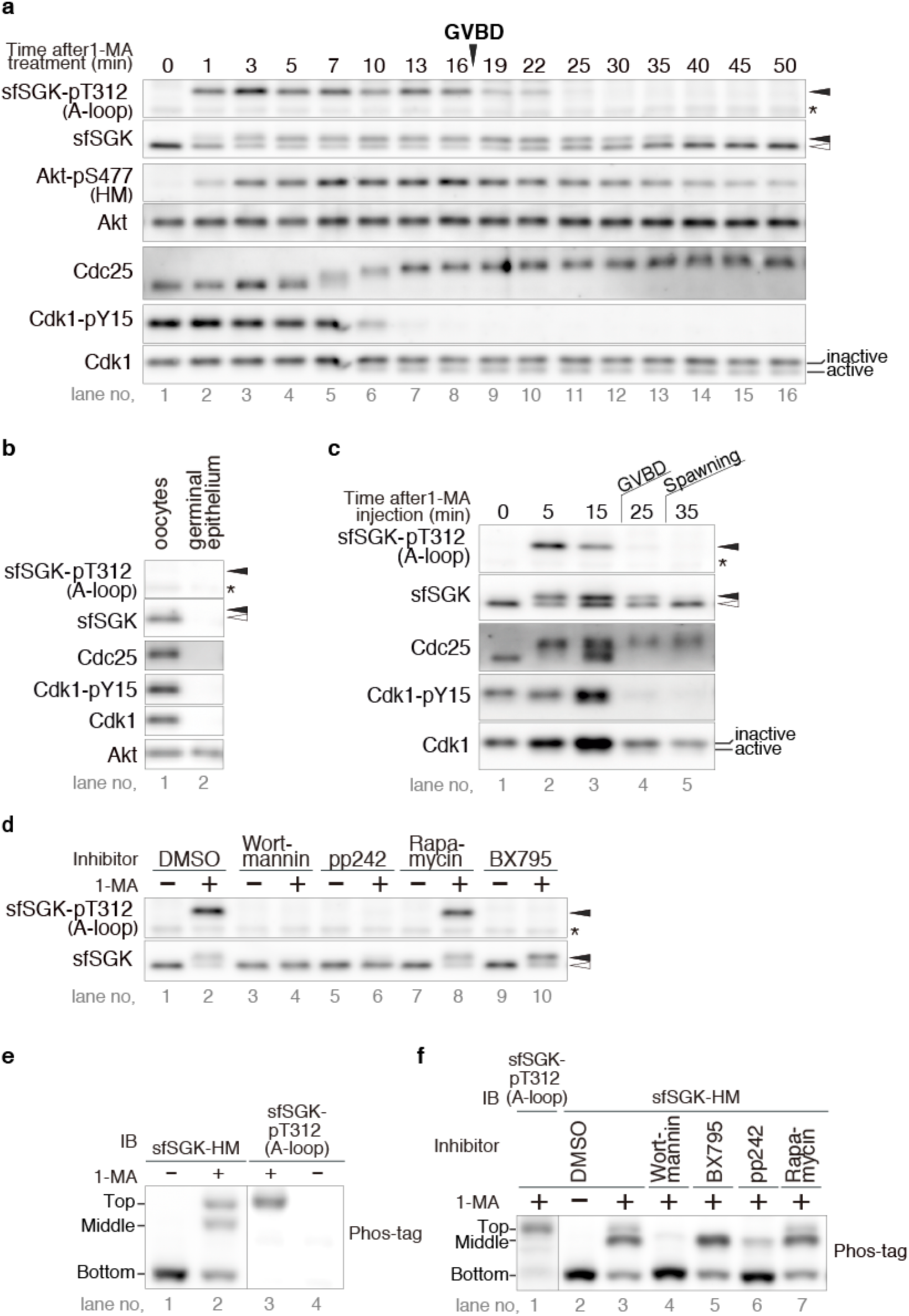
sfSGK is activated by PDK1 and TORC2 in a PI3K-dependent manner. **a** Isolated oocytes were treated with 1-MA and recovered at the indicated times, followed by immunoblotting with anti-human phospho-SGK (sfSGK-pT312 [A-loop]), anti-sfSGK-HM (sfSGK), anti-starfish phospho-Akt-HM (Akt-pS477[HM]), anti-starfish Akt C-terminal fragment (Akt), anti-starfish Cdc25 (Cdc25), anti-phospho-Cdk1 (Cdk1-pY15), and anti-PSTAIR (Cdk1) antibodies. GVBD occurred at 17 min. The result shown is representative of three independent experiments. **b** A piece of ovary was taken from female starfish, separated into oocytes and germinal epithelium, and subjected to immunoblotting with indicated antibodies. **c** To stimulate ovarian oocytes, 1-MA was injected into the body cavity of female starfish, and then two pieces of ovaries were recovered at indicated times. One piece of ovary was immediately added to sample buffer for immunoblotting with the indicated antibodies. Oocytes from the other piece of the ovary were monitored to count GVBD. GVBD occurred within 25 min, and spawning started at 30 min. The result shown is representative of two independent experiments. **d** Isolated oocytes were treated with 1-MA for 4 min in the presence of indicated inhibitors or DMSO, and then subjected to normal SDS-PAGE, followed by immunoblotting with the indicated antibodies. The result shown is representative of two independent experiments. **e** Isolated oocytes were treated with 1-MA for 4 min, and then subjected to Phos-tag SDS-PAGE, followed by immunoblotting with the indicated antibodies (IB). The result shown is representative of three independent experiments. **f** Isolated oocytes were treated with 1-MA for 4 min in the presence of indicated inhibitors, and subjected to Phos-tag SDS PAGE, followed by immunoblotting with the indicated antibodies (IB). The result shown is representative of three independent experiments. Closed and open arrowheads indicate positions of the upper and lower band of sfSGK, respectively (**a**, **b, c** and **d**). Asterisks indicate non-specific bands (**a**, **b**, **c,** and **d**).

Next, we investigated the dynamics of phosphorylation of these proteins in ovarian oocytes in the body cavity, where MI arrest occurs after GVBD^16,17,27^. To this end, we injected 1-MA into the body cavities of female starfish, and then at each time point, isolated two pieces of ovary: one for counting GVBD, and the other for immunoblotting analysis. We found that 95% of ovarian oocytes underwent GVBD within 25 min after 1-MA injection; spawning from the female started at 30 min. To prepare ovarian oocytes for immunoblotting, the piece of ovary was placed directly into sample buffer immediately after isolation. Although these samples contained not only oocytes but also other somatic cells derived from ovarian germinal epithelium, we ignored the involvement of the germinal epithelium because the sfSGK, Cdc25 and Cdk1 proteins were not detectable in the germinal epithelium (Fig, 2b). Analysis of oocyte-derived sfSGK, Cdc25, and Cdk1 proteins revealed that both sfSGK and Cdc25 in ovarian oocytes were phosphorylated within 5 min, and that Tyr15 of Cdk1 was dephosphorylated within 25 min (Fig. 2c). These results indicate that sfSGK, Cdc25, and cyclin B–Cdk1 in ovarian oocytes in the body cavity were activated similarly to those in isolated oocytes.

It should be noted that the shifted bands of sfSGK (Fig. 2a, sfSGK, lane 11-13; Fig. 2c, sfSGK, lane 4) were maintained even after disappearance of the sfSGK bands reflecting phosphorylation at the A-loop (Fig. 2a, sfSGK-pT312 [A-loop], lanes 11–13; Fig. 2c, sfSGK-pT312 [A-loop], lane 4), suggesting the existence of a phosphorylation site outside the A-loop. Given that the phosphorylation site in the HM by TORC2 is conserved in sfSGK, the mobility shift may be due to phosphorylation of the HM.

### PDK1 and TORC2 phosphorylate sfSGK in a PI3K-dependent manner

Next, we investigated the possible involvement of PDK1 and TORC2 in the activation of sfSGK in starfish oocytes. When we treated oocytes with BX795, a specific inhibitor of PDK1, A-loop phosphorylation of sfSGK after 1-MA treatment was completely blocked (Fig. 2d, lane 10), suggesting that PDK1 is required for A-loop phosphorylation. In mammalian cells, PDK1-dependent phosphorylation of the A-loop requires prior phosphorylation of the HM by mTORC2^36,38,39,41-43^. If this regulation is conserved in starfish, both the A-loop phosphorylation and HM phosphorylation of sfSGK should depend on TORC2. As expected, A-loop phosphorylation was blocked by pp242, a specific inhibitor of TOR, the catalytic subunit of TORC2 (Fig. 2d, lane 6). Notably, the mobility shift of sfSGK was reduced by pp242, but was not affected by BX795 (Fig. 2d, lanes 6 and 10), suggesting that the shift is caused by HM phosphorylation, which is dependent on TORC2 but not PDK1.

It should be noted, however, that TOR forms two types of complex: TOR complex 1 (TORC1) and TORC2^48^, both of which are inhibited by pp242. To confirm that the inhibition of sfSGK phosphorylation by pp242 was due to inhibition of TORC2 rather than TORC1, we used the TORC1 inhibitor rapamycin at a concentration previously shown to be sufficient for inhibition of TORC1 in starfish oocytes^45^. Because neither A-loop phosphorylation nor the mobility shift of sfSGK by 1-MA stimulation was blocked under this condition (Fig. 2d, lane 8), we concluded that TORC2, but not TORC1, phosphorylates sfSGK. Taken together, these observations suggest that sfSGK in starfish oocytes is activated via phosphorylation of the A-loop by PDK1 and of the HM by TORC2, and that the latter is prerequisite for the former. In mammalian cells, PDK1 and TORC2 phosphorylate SGK3 downstream of PI3K^35,36,39,43^. Consistent with this, the pan-PI3K inhibitor wortmannin blocked both A-loop phosphorylation and the mobility shift of sfSGK in starfish oocytes (Fig. 2d, lane 4), suggesting that this cascade is conserved in the 1-MA-signaling pathway.

To separate fully-activated sfSGK from sfSGK phosphorylated only on the HM, we performed Phos-tag SDS-PAGE, in which phosphorylated proteins migrate more slowly than in normal SDS-PAGE^49^. sfSGK migrated as a single band in unstimulated oocytes (Fig. 2e, the *bottom* band, lane 1). Two additional slower-migrating bands were detected after 1-MA stimulation (Fig. 2e, the *top* and *middle* bands, lane 2). Phosphorylation of the A-loop was detected only in the top band (Fig. 2e, the *top* band, lane 3), suggesting that this band corresponded to fully-activated sfSGK phosphorylated by both PDK1 and TORC2. Indeed, the top band was eliminated by the PDK1 inhibitor BX795 and the TORC1/2 inhibitor pp242 (Fig. 2f, the *top* band, lane 5 and 6), but was unchanged by the TORC1 inhibitor rapamycin. In addition, we found that the intensity of the middle band was reduced by pp242, but not by BX795 or rapamycin (Fig. 2f, *middle* band, lanes 5–7), suggesting that it represented sfSGK phosphorylated only on the HM by TORC2. Wortmannin abolished both the top and middle bands (Fig. 2f, lane 4) indicating that all phosphorylation detected using Phos-tag occurred downstream of PI3K. Taken together, these observations suggest that after 1-MA stimulus, PDK1 phosphorylates the A-loop of a subset of sfSGK that is pre-phosphorylated on the HM by TORC2 in a PI3K-dependent manner.

### Activation of sfSGK is required for rapid pH_i_ increase after 1-MA stimulus

Next, we investigated whether sfSGK is involved in the pH_i_ increase after 1-MA stimulus. Given that the anti-sfSGK-HM antibody immunoprecipitated sfSGK protein (Fig. 1c), the binding of the antibody to the HM may be strong enough to prevent the interaction of TORC2 with the HM of sfSGK and thereby block phosphorylation of the HM. Indeed, the mobility shift of sfSGK after 1-MA stimulus was blocked in oocytes injected with the antibody (Fig. 3a). More importantly, phosphorylation of the A-loop was also blocked (Fig. 3a), as observed in oocytes treated with pp242 (Fig. 2d), indicating that the anti-sfSGK-HM antibody inhibited sfSGK activation. This inhibition was specific to sfSGK, because the HM of Akt was phosphorylated even in oocytes pre-injected with the anti-sfSGK-HM antibody (Fig. 3a); accordingly, hereafter we refer to this antibody as an sfSGK-neutralizing antibody. To determine whether sfSGK activation is required for the pH_i_ increase after 1-MA treatment, we injected a pH-sensitive fluorescent dye, BCECF-dextran, into unstimulated oocytes along with the sfSGK-neutralizing antibody, and monitored the dynamics of pH_i_ by measuring fluorescence every 30 seconds before and after 1-MA addition. The sfSGK-neutralizing antibody blocked the pH_i_ increase, whereas the pH_i_ of oocytes injected with control IgG increased soon after 1-MA treatment (Fig. 3b). These results indicate that the pH_i_ increase after 1-MA treatment depends on activation of sfSGK.

**Figure 3.**
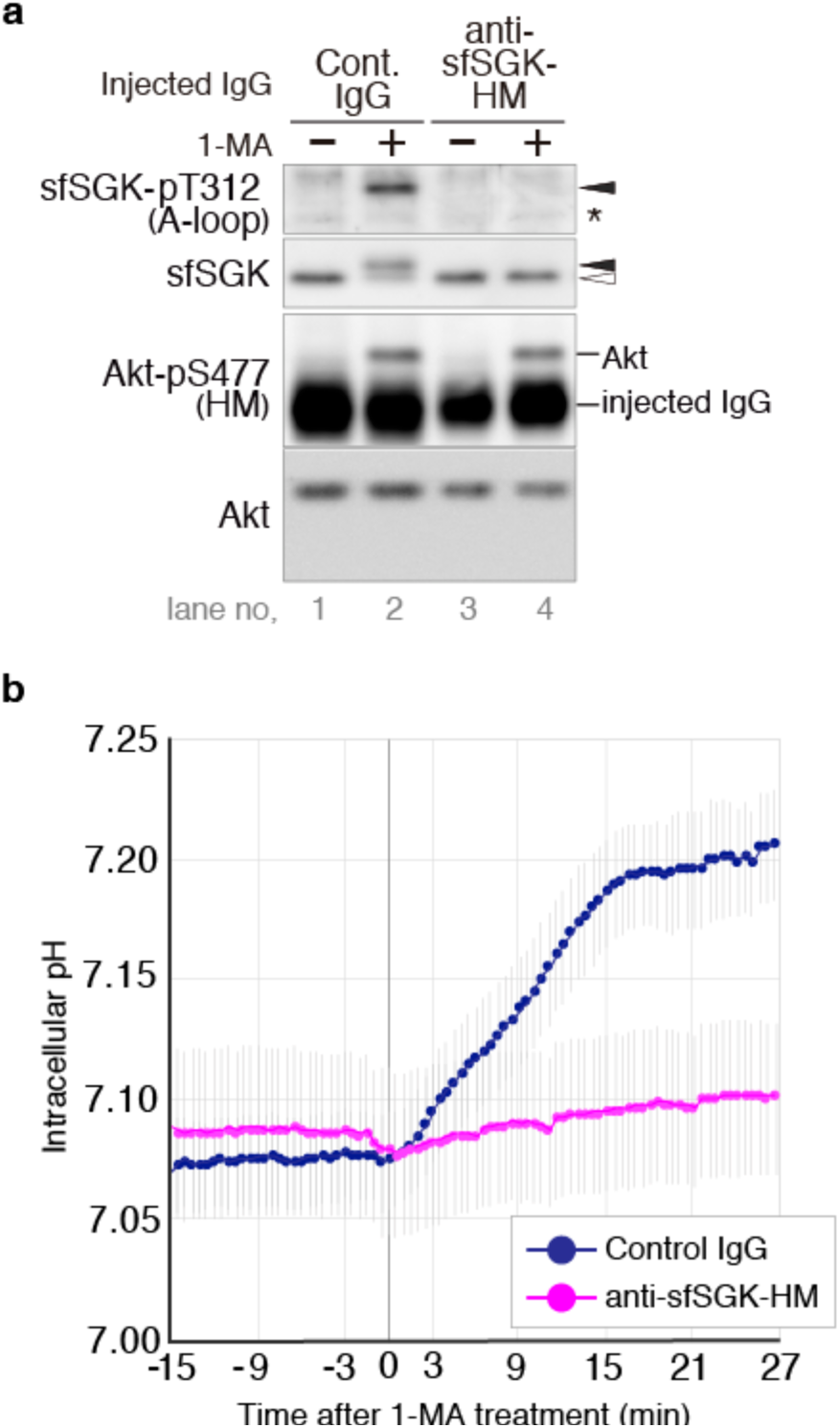
Activation of sfSGK is required for rapid pH_i_ increase after 1-MA stimulus. **a** Unstimulated oocytes were injected with an anti-sfSGK-HM antibody or control IgG, treated with 1-MA for 4 min, and then subjected to immunoblotting with indicated antibodies. Asterisk indicates non-specific bands. Closed and open arrowheads indicate positions of the upper and lower band of sfSGK, respectively. The result shown is representative of three independent experiments. **b** BCECF-dextran and either anti-sfSGK-HM antibody or control IgG were co-injected into unstimulated oocytes. After a 1-h incubation, 1-MA was added, and the fluorescence intensity ratio was measured every 30 seconds before and after 1-MA addition. pH_i_ was calculated from the fluorescence intensity ratio and plotted. Data represent means ± SE of three independent experiments.

### Activation of sfSGK, but not pH_i_ increase, is required for cyclin B-Cdk1 activation

Previously, we showed that GVBD is initiated by 1-MA in isolated oocytes even when the pH_i_ increase is blocked^16^. Nonetheless, to our surprise, GVBD was blocked by injection of the sfSGK-neutralizing antibody, whereas control IgG had no inhibitory effects on GVBD (Fig. 4a, b). When we checked the phosphorylation status of Cdc25 and Cdk1 of the oocytes injected with the sfSGK-neutralizing antibody, we found that neither hyperphosphorylation of Cdc25 nor dephosphorylation of Tyr15 of Cdk1 after 1-MA stimulus occurred (Fig, 4c), indicating that this antibody blocked 1-MA–dependent signal transduction leading to cyclin B–Cdk1 activation.

**Figure 4.**
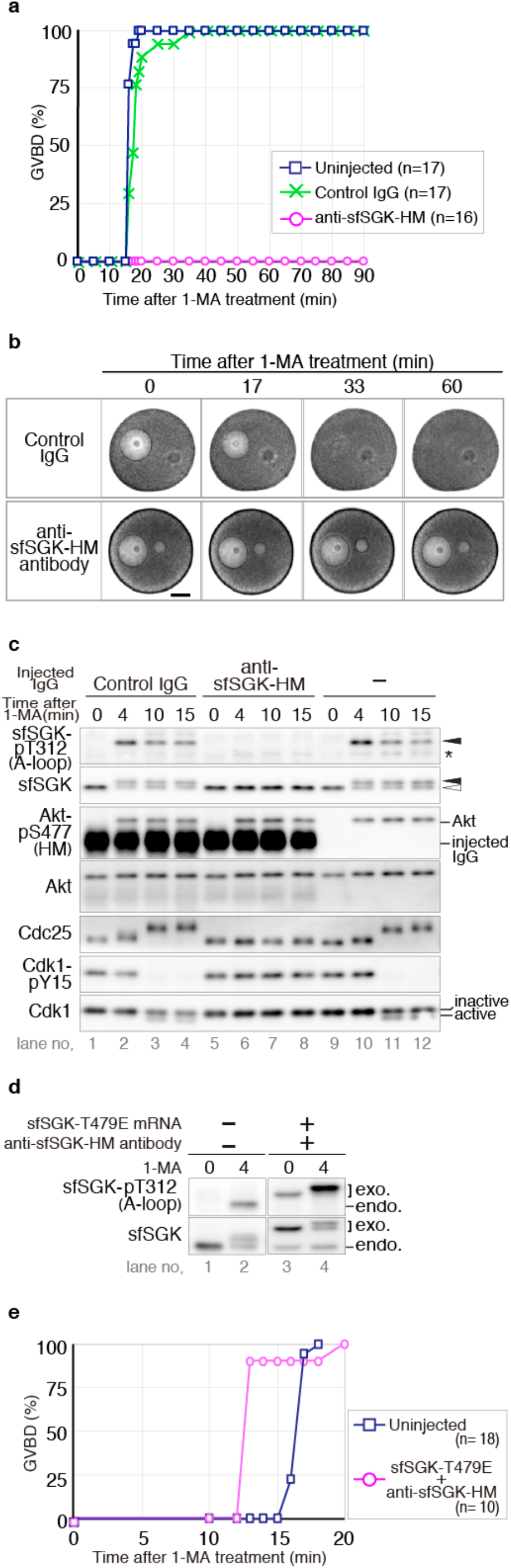
Activation of sfSGK is required for cyclin B–Cdk1 activation. **a**–**c** Unstimulated oocytes were injected with anti-sfSGK-HM antibody or control IgG, treated with 1-MA, and subjected to the following experiments: monitoring of GVBD (**a**), imaging by differential interference contrast (DIC) microscopy (**b**), and immunoblotting with the indicated antibodies (**c**) at the indicated times. “n” indicates the number of oocytes observed (**a**). An oil drop, that had been introduced along with the antibodies as a mark of the microinjection, was observed on the right of the GV (**b**). Scale bar represents 50 µm (**b**). Asterisk indicates a non-specific band (**c**). Closed and open arrowheads indicate positions of the upper band and the lower band of sfSGK, respectively (**c**). Results are representative of two independent experiments. **d**, **e** Anti-sfSGK-HM antibody was injected into unstimulated oocytes. After incubation for 1 h, mRNA encoding a mutant sfSGK (T479E) was injected into the same oocytes, followed by incubation for 17 h. As a control, uninjected oocytes were incubated for 17 h. These oocytes were treated with 1-MA, followed by immunoblotting with indicated antibodies (**d**) and counting of GVBD (**e**) at the indicated times. “exo.” and “endo.” indicate exogenous and endogenous sfSGK, respectively (**d**). “n” indicates the number of oocytes observed (**e**). Results are representative of two independent experiments.

To verify that the inhibitory effects of the antibody were caused by specific inhibition of sfSGK activation, we next performed a rescue experiment. Specifically, we replaced Thr479 of sfSGK with Glu (T479E), which was expected to mimic the negatively charged phosphate group, thereby causing PDK1-dependent A-loop phosphorylation even in the presence of the neutralizing antibody. We co-injected mRNA encoding the mutant sfSGK-T479E and the sfSGK-neutralizing antibody into unstimulated oocytes. After a 17-h incubation, the mutant was expressed and partially phosphorylated on its A-loop even in the absence of 1-MA (Fig. 4d). This partial activation did not induce GVBD. 1-MA stimulation induced enhancement of A-loop phosphorylation on the mutant sfSGK (Fig. 4d), as well as GVBD (Fig. 4e), whereas the A-loop phosphorylation and mobility shift of endogenous sfSGK were blocked in these oocytes (Fig. 4d). On the basis of these findings, we concluded that the antibody-injected oocytes are rescued by the additional expression of the mutant sfSGK-T479E, and that the inhibitory effect of the antibody is highly specific.

Interestingly, oocytes expressing the T479E mutant underwent GVBD several minutes faster than the control intact oocytes (Fig. 4e), suggesting that partial activation of the mutant in unstimulated oocytes likely shortened the time to GVBD following 1-MA treatment, further supporting the idea that sfSGK participates in 1-MA signaling leading to GVBD. We also found that the band of the T479E mutant phosphorylated on the A-loop underwent an upward mobility shift after 1-MA stimulation, indicating that the mutant was phosphorylated on a site outside the A-loop and the HM; however, the function of this phosphorylation event remains unclear. Taken together, these results suggest that activation of sfSGK is required for cyclin B–Cdk1 activation.

Although the 1-MA–induced pH_i_ increase and cyclin B–Cdk1 activation were simultaneously blocked by the sfSGK-neutralizing antibody, they were independent of each other. We base this claim on two lines of evidence: 1) The increase in pH_i_ occurred earlier than cyclin B–Cdk1 activation (Fig. 2a, 3b), and was not affected by treatment with the Cdk1 inhibitor, roscovitine (Supplementary Fig. 1), indicating that the pH_i_ increase is independent of cyclin B–Cdk1 activation. 2) GVBD in isolated oocytes, in which initial pH_i_ before 1-MA stimulation is ~7.0, is initiated even when the pH_i_ increase was blocked by an NHE inhibitor or by treatment with sodium-free ASW^16^, indicating that the cyclin B–Cdk1 activation leading to GVBD is independent of the pH_i_ increase (see also below, Fig. 5a). Thus, the pH_i_ increase and cyclin B–Cdk1 activation are mutually independent events.

**Figure 5.**
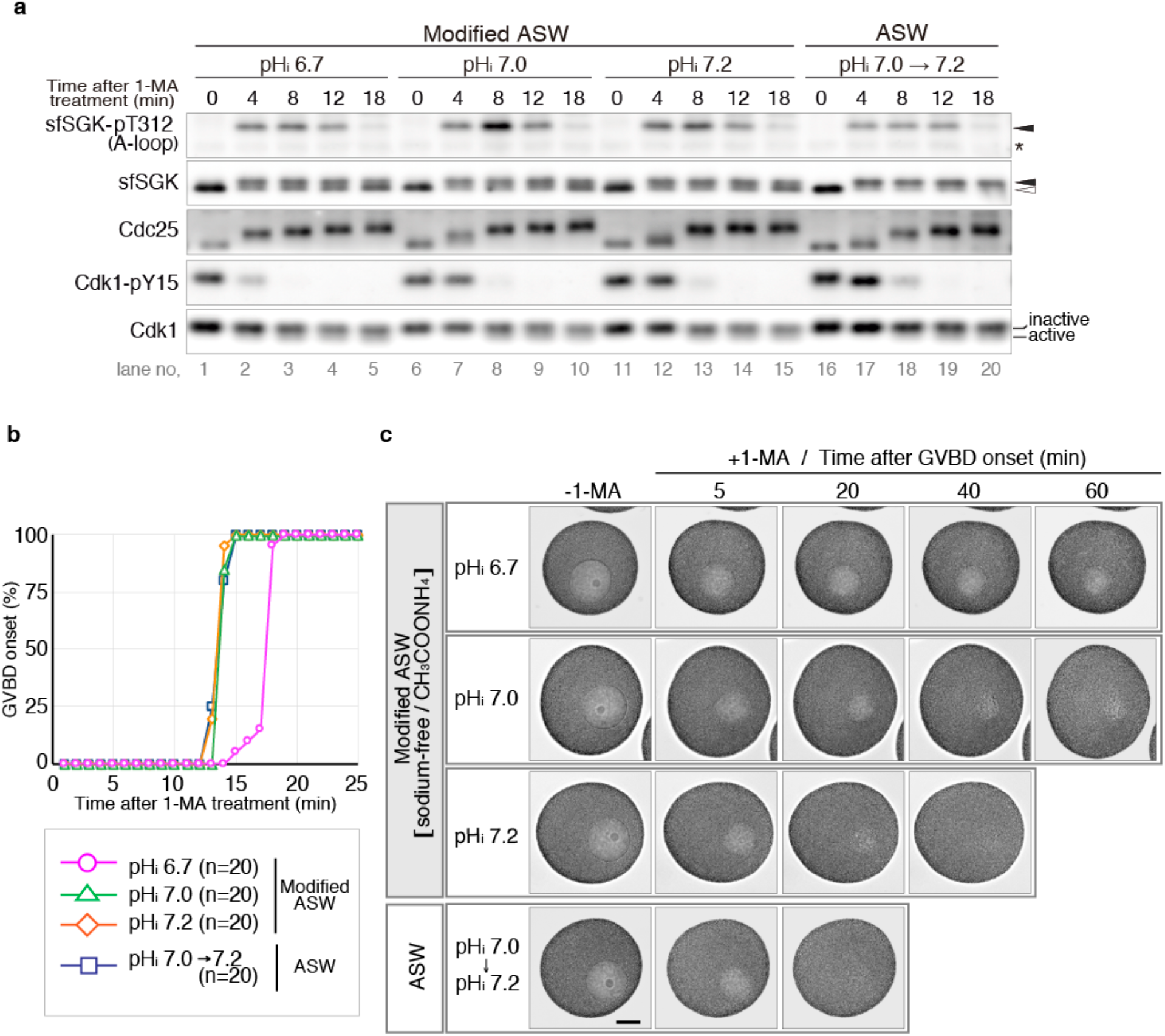
Reduced pH_i_ delays onset of GVBD and blocks completion of GVBD without affecting cyclin B–Cdk1 activation. **a** To clamp pH_i_ at 6.7, 7.0, and 7.2, unstimulated oocytes were incubated with sodium-free artificial seawater containing CH_3_COONH_4_ (Modified ASW) for 20 min. As a control, unstimulated oocyteswere incubated in artificial seawater (ASW) for 20 min, in which pH_i_ is approximately 7.0, andincreases to ~7.2 after 1-MA treatment^17^ (see also control IgG–injected oocytes in Fig. 3b). These oocytes were treated with 1-MA and analyzed by immunoblotting with indicated antibodies. Asterisk indicates a non-specific band. Closed and open arrowheads indicate the positions of the upper and lower bands of sfSGK, respectively. **b** Oocytes treated as described in **a** were monitored to count onset of GVBD. “n” indicates the number of oocytes observed. Morphology of GVBD onset is shown in Supplementary Fig. 2. **c** Time-lapse DIC images of oocytes treated as described in **a** were taken every 10 seconds before and after addition of 1-MA. Images at selected times after GVBD onset are shown. Scale bar represents 50 µm. For complete image sequences, see Supplementary movies 1–4. For Z-stack images acquired after the time-lapse imaging, see Supplementary Fig. 3. Results in all panels are representative of two independent experiments.

### Reduced pH_i_ delays onset of GVBD and blocks completion of GVBD without affecting cyclin B–Cdk1 activation

To elucidate the role of the pH_i_ increase, we investigated effect of altered pH_i_ on 1-MA–induced meiotic resumption in starfish oocytes. To clamp pH_i_, oocytes were incubated in sodium-free ASW containing CH_3_COONH_4_ (modified ASW), in which the pH_i_ was adjusted to the desired value with CH_3_COONH_4_ but did not increase after 1-MA stimulation because of the absence of sodium ion^17^ (see also Materials & Methods). We clamped pH_i_ at 6.7, corresponding to the value in unstimulated ovarian oocytes^17^; as well as 7.0 and 7.2, similar to the values in isolated oocytes before and after 1-MA stimulus, respectively^17^. First, we examined effects of pH_i_ clamping on sfSGK and cyclin B–Cdk1 activation after 1-MA stimulus. Immunoblotting analysis showed that the phosphorylation status of sfSGK, Cdc25, and Cdk1 in oocytes at all clamped pH_i_ values was basically same as in oocytes in ASW (Fig. 5a). This observation suggests that 1-MA–induced signaling leading to cyclin B–Cdk1 activation does not depend on pH_i_, and further supports our conclusion that the pH_i_ increase is not required for cyclin B–Cdk1 activation.

After cyclin B–Cdk1 activation, the onset of GVBD was delayed for several minutes at a clamped pHi of 6.7 (Fig. 5b), as previously reported^17^. At clamped pH_i_ values of 7.0 and 7.2, the times of GVBD onset were similar to those in ASW (Fig. 5b). At all clamped pH_i_ values, the morphology of GVBD onset was indistinguishable from those in ASW: before 1-MA stimulation, the rim of GV looked like a clear line under the microscope; at GVBD onset, cytoplasmic granules started to invade from the rim into the inner GV area (Supplementary Fig. 2). These results suggest that after cyclin B–Cdk1 activation, time to onset of GVBD is weakly pH_i_-dependent.

More importantly, we observed a remarkable defect after onset of GVBD. In ASW, cytoplasmic granules occupied the whole inner GV area within 15 min after GVBD onset, indicating completion of GVBD (Fig. 5c, Supplementary Movie 1, Supplementary Fig. 3). At a clamped pH_i_ of 6.7, however, we found that the invasion of granules proceeded slowly and stalled around 40 min, remaining incomplete even 60 min after GVBD onset (Fig. 5c, Supplementary Movie 2, Supplementary Fig. 3). At a clamped pH_i_ of 7.0, invasion proceeded further and finished with a trace of inner GV area within 40–50 min (Fig. 5c, Supplementary Movie 3, Supplementary Fig. 3). At clamped pH_i_ of 7.2, the granules invaded and ultimately distributed homogenously throughout the oocytes within 20–30 min after GVBD onset (Fig. 5c, Supplementary Movie 4, Supplementary Fig. 3). These observations suggest that progression of granule invasion is sensitive to pH_i_; in particular, completion of GVBD is drastically impaired at a pH_i_ of 6.7. Given that the pH_i_ of unstimulated ovarian oocytes is around 6.7 (ref. ^17^), the pH_i_ increase mediated by sfSGK is likely required for completion of GVBD in ovarian oocytes.

## Discussion

The results of this study show that sfSGK is activated by TORC2 and PDK1 in a PI3K-dependent manner after 1-MA stimulus in starfish oocytes. We found that sfSGK is required for regulation of two mutually independent pathways leading to pH_i_ increase and cyclin B–Cdk1 activation. Furthermore, we identified the presence of a pH_i_-sensitive process in meiotic resumption: although 1-MA signaling leading to cyclin B–Cdk1 activation is pH_i_-independent, the subsequent onset time and completion of GVBD are pH_i_-dependent.

On the basis of previous studies and the present findings, we propose a model for meiotic resumption in ovarian oocytes (Fig. 6). In this model, sfSGK exerts two essential functions: cyclin B–Cdk1 activation, which induces meiotic resumption, and pH_i_ increase, which promotes completion of GVBD (Fig. 6). In living starfish, ovarian oocytes reside in coelomic fluid in the body cavity. Previously, we reported that the coelomic fluid has relatively high CO_2_ and low O_2_ concentrations relative to ASW^17^. Under these gas conditions, the pH_i_ of unstimulated ovarian oocytes is around 6.7, and increases after 1-MA stimulation^17^. As shown in Fig. 2c, sfSGK is activated in ovarian oocytes after 1-MA stimulus in the body cavity. Subsequently, sfSGK likely induces the pH_i_ increase and cyclin B–Cdk1 activation in ovarian oocytes (Fig. 6), as demonstrated in isolated oocytes (Figs. 3 and 4). Thereafter, cyclin B–Cdk1 causes GVBD. However, if pH_i_ in ovarian oocytes was not increased from 6.7 soon after 1-MA stimulus, GVBD would initiate after a delay and fail to complete, as we observed when pH_i_ was clamped at 6.7 in isolated oocytes (Fig. 5). Thus, in ovarian oocytes, the sfSGK-dependent pH_i_ increase would determine the time of GVBD onset after cyclin B–Cdk1 activation and be a prerequisite for completion of GVBD (Fig. 6).

**Figure 6.**
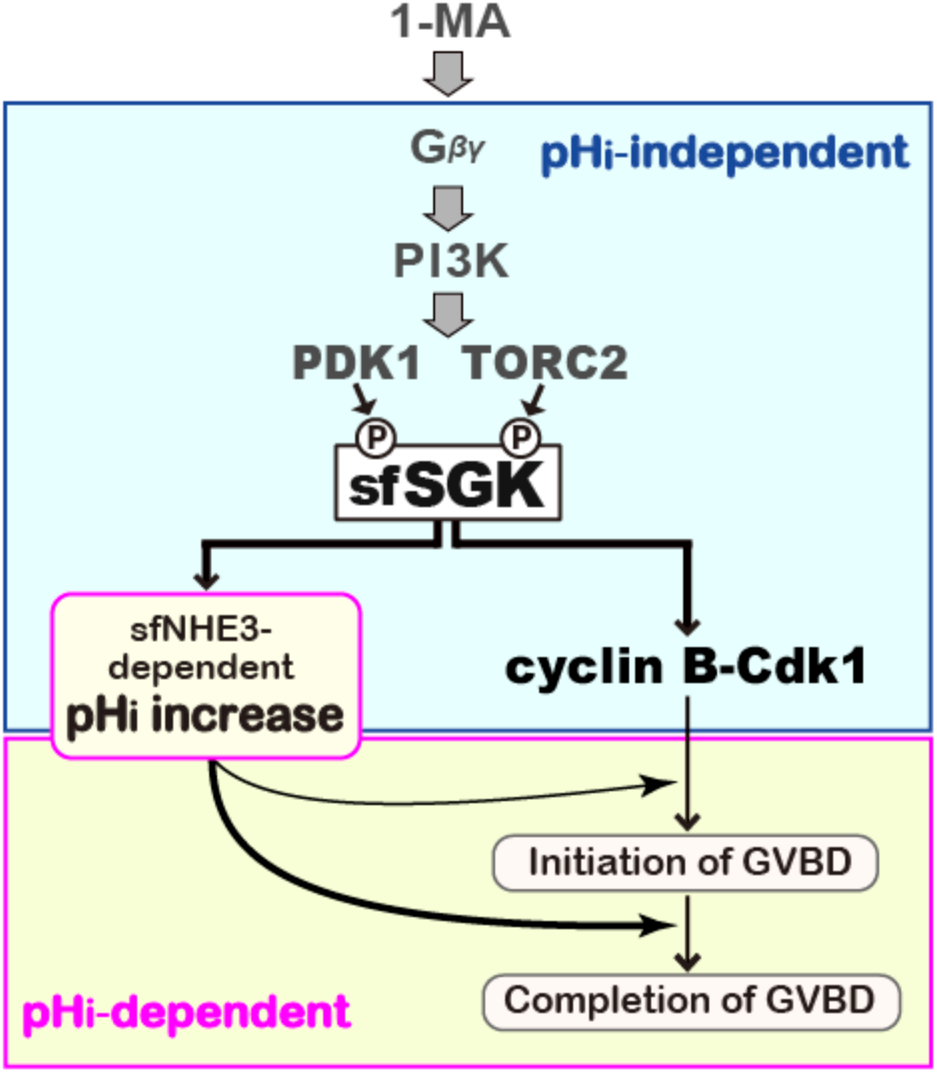
Model for sfSGK-dependent meiotic resumption: sfSGK induces cyclin B–Cdk1 activation to resume meiosis and pH_i_ increase to complete GVBD in ovarian oocytes. A model for 1-MA–induced meiotic resumption in ovarian oocytes was prepared on the basis of previous and current findings. 1-MA stimulus induces the release of Gβγ from Gαi, leading to PI3K activation. In a manner that depends on PI3K, sfSGK is activated via phosphorylation of the A-loop by PDK1 after prior phosphorylation of the HM by TORC2. Activation of sfSGK is essential for cyclin B–Cdk1 activation and sfNHE3-dependent pH_i_ increase. These SGK-dependent pathways are pH_i_-independent (blue box). On the other hand, time to GVBD onset after cyclin B–Cdk1 activation and completion of GVBD are pH_i_-dependent (yellow box), and both are defective at pH_i_ of 6.7.Because pH_i_ of ovarian oocytes before 1-MA stimulation in body cavity is ~6.7, the sfSGK-dependent pH_i_ increase after 1-MA stimulus accelerates GVBD onset and is required for completion of GVBD in ovarian oocytes. By contrast, the pH_i_ of isolated oocytes before 1-MA stimulation in ASW is ~7.0, which is already permissive for GVBD. Therefore, a pH_i_ increase is not essential for GVBD in isolated oocytes.

In the ovaries, starfish oocytes arrest at MI after GVBD ^16,17,27^. This arrest is released upon spawning, at which time the oocytes are ready to be fertilized^16,17,27^. Fertilization during the period between release from the MI arrest and the first polar body formation is important for monospermy, because insemination before GVBD or after polar body formation tends to result in polyspermy^27,50^. Thus, the sfSGK-dependent pH_i_ increase soon after 1-MA may ensure that ovarian oocytes proceed to MI arrest before spawning, helping to promote monospermy. In addition, to ensure fertilization before polar body formation, MI arrest must be maintained until spawning^27^. To maintain MI arrest, sfSGK-dependent pH_i_ increase should not exceed 7.0, because release from MI arrest occurs at pH_i_>7.0 (ref. ^16,17,26,27^). In this study, A-loop phosphorylation of sfSGK became undetectable around the time of the end of pH_i_ increase (Fig. 2a, c). Therefore, sfSGK inactivation may be important to stop the pH_i_ increase for maintaining MI arrest. Interestingly, the A-loop was dephosphorylated even in the presence of phosphorylated HM (Fig. 2a, c), implying the presence of regulatory mechanism of A-loop dephosphorylation that is not associated with dephosphorylation of the HM, such as up-regulation of an A-loop phosphatase. To our knowledge, regulation of SGK inactivation has never been investigated in any animal, and this issue should be examined in a future study.

Although mammalian SGKs are involved in cell proliferation^51,52^, their function in M-phase remains unclear. Here, we showed that sfSGK is required for cyclin B–Cdk1 activation in starfish oocytes (Fig. 4). In parallel with the present study, Hiraoka et al. demonstrated that sfSGK directly phosphorylates Cdc25 and Myt1 to activate cyclin B–Cdk1, and that sfSGK, rather than Akt, is the major kinase responsible for induction of cyclin B–Cdk1 activation^53^. These are the first reports to demonstrate the involvement of SGK in cyclin B–Cdk1 activation.

In addition, we showed that sfSGK is required for the pH_i_ increase (Fig. 3b). Previously, we reported that this increase depends on sfNHE3^16,17,25^. In human cultured cells, NHE-dependent pH_i_ increase is upregulated by phosphorylation of C-terminal region of NHE that leads to NHE activation^7,9^, and/or by accumulation of NHE protein on the plasma membrane through trafficking of endosomal NHE to the plasma membrane^6^. Although it is not known whether these mechanisms are functional in starfish oocytes, it is clear from our previous work that the C-terminus of sfNHE3 is phosphorylated by human SGK1 *in vitro*^25^. Therefore, we speculate that sfSGK upregulates sfNHE3 by direct phosphorylation of its C-terminus.

In the process of GVBD in starfish oocytes, the nuclear pore complex on nuclear envelopes (NEs) of the GV are first disassembled^54-56^, and then the NEs are rapidly broken into membrane fragments, followed by complete mixing of cytoplasm and nucleoplasm^54,55^. GVBD onset, as judged by differential interference contrast (DIC) microscopy in this study, likely represents the time of membrane fragmentation, as a clear boundary between cytoplasm and GV was still observed by DIC during nuclear pore disassembly^55^. One possible interpretation of the remarkable blockage of GVBD completion at a clamped pH_i_ of 6.7 (Fig. 5b, c) is that incomplete NE fragmentation occurred and disrupted invasion of cytoplasmic granules. Given that NE fragmentation depends on F-actin shell formed on the inner surface of the GV^56^, our present observations imply that the formation or function of this shell is blocked at pH_i_ 6.7. Although this possibility is yet to be tested, F-actin architecture is known to be disrupted by lower pH_i_, at least in mouse mammary tissue^57^. pH_i_ increase after release from ProI arrest has also been reported in frog^13^, urodele^58^, and surf clam^12^. In *Xenopus* oocytes, NHE-dependent pH_i_ increase after hormonal stimulation^13,14,59^ has been suggested to accelerate cyclin B–Cdk1 activation by promoting accumulation of Mos protein, which participates in cyclin B–Cdk1 activation^60^. By contrast, we found that cyclin B–Cdk1 activation was independent of the pH_i_ increase. One potential reason for this difference could be cyclin B–Cdk1 activation in starfish oocytes does not require synthesis of new protein^61^, including Mos^62^. In addition, pH_i_ increase in *Xenopus* oocytes plays a role in migration of GV to the animal pole before GVBD^63^. Such migration does not occur in starfish oocytes because the GV already resides at the animal pole even in ProI-arrested oocytes. Instead, we found that completion of GVBD depends on pHi. Thus, starfish and *Xenopus* oocytes provide suitable models for understanding different aspects of pH_i_-dependent regulation of the meiotic cell cycles.

In summary, we showed that sfSGK is required for pH_i_ increase and cyclin B–Cdk1 activation in starfish oocytes, and that completion of GVBD is a pH_i_-sensitive process. These findings reveal a novel role for SGK in control of meiotic cell cycles and emphasize the importance of tight regulation of pH_i_ in oocyte meiosis.

## Materials and methods

### Oocyte preparation

Starfish (*Asterina pectinifera*) were collected on Pacific coast of Japan, and kept in laboratory aquaria supplied with circulating seawater at 14°C. Fully-grown ProI-arrested oocytes were isolated from ovaries and treated with calcium-free artificial seawater (calcium-free ASW: 476 mM NaCl, 10 mM KCl, 36 mM MgCl_2_, 18 mM MgSO_4_, 20 mM H_3_BO_3_, pH 8.2) to remove follicle cells. Subsequently, the oocytes were kept in artificial seawater (ASW: 462 mM NaCl, 10 mM CaCl_2_, 10 mM KCl, 36 mM MgCl_2_, 18 mM MgSO_4_, 20 mM H_3_BO_3_, pH 8.2) until use. For 1-MA stimulation, the isolated oocytes were incubated with ASW containing 0.5 µM 1-MA at 23°C, and then collected for analysis. To induce meiotic resumption of ovarian oocytes in the body cavity, 1 ml of calcium-free ASW without boric acid (462 mM NaCl, 10 mM KCl, 36 mM MgCl_2_, 17 mM MgSO_4_) containing 0.5 mM 1-MA was injected into the body cavity of female starfish. Pieces of ovary containing stimulated oocytes were directly recovered from the body cavity, and then immediately added into a sample buffer (0.25 M Tris-HCL [pH 6.8], 40% glycerol, 8% SDS, 20% 2-mercaptoethanol, 0.01% bromophenol blue). To compare proteins in germinal epithelium with those in oocytes, a piece of ovary was isolated from the body cavity and separated into germinal epithelium and oocytes. These fractions were separately recovered into different tubes, each of which contained an equal volume of sample buffer, boiled at 95°C for 5 min, and subjected to immunoblotting analysis.

### cDNA cloning of starfish SGK

To isolate a cDNA encoding the starfish homolog of SGK, total mRNA was isolated from ProI-arrested ovarian oocytes using RNAwiz (Ambion, Thermo Fisher Scientific; Waltham, MA, USA). The first-strand cDNA library was prepared from total mRNA using the TAKARA RNA PCR Kit (AMV) Ver.2.1 and Prime Script Reverse Transcriptase (Takara Bio; Shiga, JP) with random 9-mers. A 250-bp cDNA fragment (fragment 1) was first obtained by reverse-transcriptase (RT)-PCR with degenerate primers designed based on amino acid sequences 18 conserved among human, mouse, sea urchin, and *C. elegans* SGKs (forward: YAVKVL, 5’-TAYGCNGTNAARGTNYT-3’, and reverse: FYAAEIA, 5’-AARATRCGNCGNCTYTCDCG-3’). Because 3’ and 5’ rapid amplification of cDNA ends (RACE) with specific primers designed against fragment 1 were unsuccessful, we obtained another fragment (fragment 2) that partially overlapped with fragment 1, as follows: a forward specific primer (5’-CCCGACAAGCTCTAC-3’) was designed against fragment 1, and a reverse degenerate primer (GLPPFY, 5’-CCNRANGGNGGNAARAT-3’) was designed against a conserved amino acid sequence in the SGK alignment that is located in a region approximately 85 amino acids C-terminal to the region of the first reverse degenerate primer. The 360-bp fragment 2 was obtained by RT-PCR using these primers. Next, to identify the 3’ end, another first-strand cDNA library was generated from the total mRNA using RNA PCR Kit (AMV) Ver. 2.1 and Prime Script Reverse Transcriptase (Both, Takara Bio; Shiga, JP) using the oligo dT-Adaptor primer from the kit. Then, RT-PCR was performed on the cDNA library using a specific forward primer designed against the 5’ region of fragment 2 (5’-CCCCAGAAGTCTTGAAGAA-3’) and the adaptor primer (5’-GTTTTCCCAGTCACGAC-3’). Using the RT-PCR product as a template, nested PCR was performed with another forward specific primer (5’-GGAGTATGATCGTAGTGTAG-3’) and the adaptor primer, resulting in amplification of a ~600 bp 3’ RACE product. Next, the 5’ ends were identified by RACE using 5’-Full RACE Core Set (Takara Bio; Shiga, JP) with specific primers (reverse for generating first-strand cDNA: 5’-CCAAAGTCTGTCAAC-3’; forward for first PCR: 5’-CAGAGATTTGAAGCCGGAG-3’, reverse for first PCR: 5’-GTTCAGGGAAGCATCTCTTCT-3’; forward for nested PCR: 5’-GCCGGAGAACATTTTGATTG-3’, reverse for nested PCR: 5’CTCTCTCTGCAGATGGAAGA-3’), resulting in amplification of a 1.5-kb 5’ RACE product. The full-length ORF was PCR-amplified from the cDNA library generated using random 9mers with pfx50 DNA polymerase (Thermo Fisher Scientific) and specific primers (forward: 5’-GGGGCCCCTGGGATCCATGGCTGCCAACAGTACTAG-3’; reverse: 5’-GTCGACCCGGGAATTCTTAGTCGTCAAATGCCTCG-3’). The amplified cDNA was inserted into vector pGEX-6P-3 (GE Healthcare; Little Chalfont, UK) using the In-Fusion HD Cloning Kit (Takara Bio; Shiga, JP) and sequenced.

### Immunoprecipitation

For preparation of oocyte extracts, oocytes were recovered in ASW before and after 1-MA treatment. After freezing in liquid N_2_, lysis buffer [80 mM β-glycerophosphate, 20 mM EGTA, 10 mM Mops pH 7.0, 100 mM sucrose, 100 mM KCl, 1 mM DTT, 0.5 % NP-40, 1% protease inhibitor cocktail (Nacalai Tesque; Kyoto, JP), and 1% phosphatase inhibitor cocktails 1 and 2 (Sigma; St Louis, MO, USA)] was added to the oocytes, followed by vortexing and centrifugation at 12,500′ *g* for 15 min. The supernatant was used as the extract. For immunoprecipitation, 15 µl of oocyte extract (1 oocyte per 1 µl) was added to 3 µl of protein A–Sepharose 4B (Sigma) to which 7.0 µg of anti-sfSGK-HM antibody or control IgG had been pre-adsorbed and cross-linked using borate buffer containing 25 mM DSS (25 mM DSS, 0.2 M H_3_BO_3_ [pH 9.0], 0.2 M NaOH), followed by incubation for 90 min on ice. Beads and flow-through were separated, and sample buffer was added. The samples were boiled at 95°C for 5 min and subjected to immunoblotting analysis.

### Chemicals

Wortmannin (LC laboratories; Woburn, MA, USA), pp242 (Cayman Chemical; Ann Arbor, MI, USA), rapamycin (FUJIFILM Wako pure chemical; Osaka, JP), BX795 (Enzo Life Sciences; New York, NY, USA), and roscovitine (Merck, Darmstadt, BRD) were dissolved in DMSO at 20 mM as a stock solution, and used at final concentrations of 40, 2, 20, 6, and 30 µM, respectively.

### DNA construct

A mutant sfSGK in which Thr479 was replaced with Glu was generated by PCR using PrimeSTAR MAX Polymerase (Takara Bio; Shiga, JP) with primers containing the point mutation (forward: 5’-GGCTTCGAATATGTCCCACCAAGCGAG-3’, and reverse: 5’-GACATATTCGAAGCCAGTAAAGGCAGAG-3’). The mutated cDNA was inserted into the pSP64-S-based vector, in which its SP6 promoter was replaced with a T7 promoter^32^, with the *A. pectinifera* cyclin B Kozak sequence (5’-TACAAT-3’) and a 3′ FLAG-tag at C-terminus.

### Antibodies

The anti-sfSGK-ΔN50 rabbit polyclonal antibody was raised against a Nus- and His-tagged recombinant sfSGK fragment lacking the N-terminal 50 amino acids including the PX domain. The antibody was purified from the antiserum using PVDF membrane on which a recombinant GST-tagged sfSGK protein had been blotted. The anti-sfSGK-HM rabbit polyclonal antibody was raised against a synthetic peptide representing the 17 C-terminal amino acids containing the HM (Ser473-Asp489) of sfSGK, and affinity-purified using antigen peptide–conjugated Sulfo Link Coupling Resin (Thermo Fisher Scientific). The control IgG was purified from rabbit pre-immune serum using protein A–Sepharose 4B (Sigma). For microinjection, the anti-sfSGK-HM antibody and control IgG were further concentrated using a 50-kDa cut-off Amicon Ultra (Merck; Rahway, NJ, USA), and the buffer was replaced with injection buffer (0.05% NP-40 in PBS buffer, pH 7.2). Preparation of anti-starfish phospho-Akt (Ser477) rabbit polyclonal antibody^45^, anti-starfish Akt (sfAkt) C-terminus rabbit polyclonal antibody^46^, anti-starfish Cdc25 (sfCdc25) rabbit polyclonal antibody^47^, and anti-PSTAIR antibody for Cdk1 (ref. ^45^) were described previously. The anti-phospho-Tyr15 Cdk1 antibody and anti-human phospho-SGK rabbit polyclonal antibody [p-SGK (Thr 256)-R] were purchased from Cell Signaling Technology (Danvers, MA, USA) and Santa Cruz Biotechnology (Dallas, TX, USA), respectively. Peroxidase-conjugated anti-rabbit IgG and anti-mouse IgG, used as secondary antibodies in immunoblotting, were purchased from GE healthcare (Little Chalfont, UK) and Dako (Glostrup, Denmark), respectively.

### Microinjection

Microinjection was performed as previously described^64,65^. The anti-sfSGK-HM antibody or control IgG was injected into unstimulated oocytes at 23°C at a final concentration in the oocyte of 65.2 µg/ml, followed by incubation for 1 h. For expression of sfSGK (T479E), the mutant was transcribed from the pSP64-S-based vector construct using the mMESSAGE mMACHINE kit (Ambion, Thermo Fisher Scientific), dissolved in water, and injected into unstimulated oocytes (30 pg per oocyte). After further incubation for 17 h at 20°C, these oocytes were treated with 0.5 µM 1-MA and recovered for immunoblotting analysis at 4 min. The rest of the oocytes were observed to count GVBD.

### Immunoblotting

Oocytes (five for normal SDS-PAGE, six for Phos-tag-SDS-PAGE) were recovered in 3 µl of seawater, added directly to 3 µl of 2′ sample buffer, and immediately frozen in liquid N_2_. After thawing and boiling for 5 min at 95°C, proteins were separated on polyacrylamide gels (8% for sfSGK only, 8.5% for sfSGK and Cdk1). For Phos-tag SDS-PAGE, 8% polyacrylamide gel containing 60 µM Phos-tag acrylamide (FUJIFILM Wako Pure Chemical co.; Osaka, JP) and 600 µM ZnCl_2_ was used. Immunoblotting was performed as previously described^32^. Antibodies used for immunoblotting were as follows: anti-sfSGK-ΔN50 (1:2,000, in TBS-T), anti-sfSGK-HM (1:1000, in TBS-T), anti-human phospho-SGK (1:400, in TBS-T), anti-sfAkt-pS477 (1:600, in Can Get Signal Immunoreaction Enhancer Solution 1 [Can Get sol. 1], TOYOBO, Osaka, Japan), anti-sfAkt-C-terminus (1:400, in Can Get sol. 1), anti-sfCdc25 (1:2000, in Can Get sol. 1), an anti-Cdk1-pY15 (1:1000, in Can Get sol. 1), and anti-PSTAIR (1:50,000, in Can Get sol. 1). horseradish peroxidase (HRP)-conjugated anti-rabbit IgG (1: 2000, in TBS-T or Can Get sol. 2) and anti-mouse IgG (1:2000, in Can Get sol. 2) were used as secondary antibodies. Proteins reacting with the antibodies were visualized with ECL prime (GE healthcare; Little Chalfont, UK), and digital images were acquired on a LAS4000 mini imager (Fujifilm; Tokyo, JP).

### Measurement of pH_i_ with BCECF-dextran

A dextran (10-kDa)-conjugate of 2’,7’-bis-(2-carboxyethyl)-5(and-6)-carboxyfluorescein (BCECF) (Molecular Probes, Thermo Fisher Scientific) was dissolved at a concentration of 2 mM in injection buffer (100 mM potassium aspartate, 10 mM HEPES, 0.05% NP-40, pH 7.2). BCECF-dextran (20 pl per oocyte) and either the sfSGK-HM antibody or control IgG (final 65.2 µg /ml in an oocyte) were co-injected into unstimulated oocytes. After incubation for 1 h, the pH_i_ of these oocytes was measured every 30 seconds before and after 1-MA treatment. pH_i_ measurements with BCECF-dextran were made by determining the pH-dependent ratio of its emission intensity (detected at 535 nm) when excited at 495 nm versus its isosbestic point of 436 nm. Calibration of pH_i_ from the intensity ratio was performed as previously described^17^. All fluorescence images were captured and analyzed using a NIKON ECLIPSE Ti-U fluorescent microscope (Nikon Instech; Tokyo, JP), which was connected to a CMOS camera (ORCA-Flash2.8, C11440-10C; Hamamatsu Photonics; Hamamatsu, JP) controlled by an HCImage processing system (Hamamatsu Photonics; Hamamatsu, JP). Excitation light from a xenon lamp alternated between 436 nm and 495 nm under computational control. The emitted light passed through a dichroic beam splitter at 505 nm and through a 510–560 nm emission filter.

### Clamping pH_i_ of oocytes

NH_4_Cl dissolved in seawater forms NH_3_, which crosses cell membrane easily and binds H^+^, causing an increase in pH_i_. By contrast, when CH_3_COONa is dissolved in seawater, it forms CH_3_COOH, which crosses cell membrane easily and releases H^+^ into the cell, causing pHi to decrease. By applying this principle, Hamaguchi *et al.* found that extracellular pH (pH_o_) and pH_i_ were almost equal when sea urchin eggs were treated with ASW containing 20 mM CH_3_COONa and NH_4_Cl^66^. We showed previously that when starfish oocytes are incubated in sodium-free ASW containing CH_3_COONH_4_ at various pH values (modified ASW: 480 mM choline chloride, 55 mM MgCl_2,_ 5 mM KCl, 10 mM PIPES, 10 mM HEPES, 20 mM CH_3_COONH_4_, 9.2 mM CaCl_2_), the pH_i_ of oocytes was invariably ~0.2 units higher than pH_o_^17^. Hence, to clamp pH_i_ of oocytes at 6.7, 7.0, and 7.2, unstimulated oocytes were incubated with modified ASW with a pH of 6.5, 6.8, and 7.0, respectively, for 20 min. These oocytes were then treated with 1-MA, followed by observation of GVBD or immunoblotting analysis. It should be noted that because sodium ion is not present in modified ASW, the sfNHE3-dependent pH_i_ increase mediated by Na^+^/H^+^ exchange does not occur, so pHi is maintained at the clamped value even after 1-MA treatment. As a control, isolated oocytes were incubated in ASW (pH_o_ 8.2), in which the pHi of unstimulated oocytes was ~7.0. Because ASW contains sodium ions, a sfNHE3-dependent pH_i_ increase to 7.2–7.3 occurs after 1-MA treatment^17^; see also the control IgG–injected oocytes in Fig. 3b.

### Oocyte observation

GVBD onset was visually judged by focusing the microscope on the equatorial plane of the GV: the rim of intact GV appears as a sharp line in unstimulated oocytes, but becomes fuzzy at GVBD onset. All DIC images and DIC image sequences were captured on a NIKON ECLIPSE Ti-U microscope (Nikon Instech; Tokyo, JP), which was connected to a CMOS camera (ORCA-Flash2.8, C11440-10C; Hamamatsu Photonics; Hamamatsu, JP) controlled by a HCImage processing system (Hamamatsu Photonics; Hamamatsu, JP). For time-lapse recording, DIC images were collected every 10 seconds. Z-stack images were captured soon after time-lapse imaging.

### Data availability

Data supporting the findings of this study are available within the article or upon reasonable request to the corresponding authors.

## Supporting information

Supplementary Movie 1_ASW

Supplementary Movie 2_pHi6.7

Supplementary Movie 3_pHi7.0

Supplementary Movie 4_pHi7.2

## Acknowledgements

We thank M. Matsushita for raising the anti-starfish SGK-ΔN50 antibody, M. Terasaki for providing vector pSP64-S, and E. Okumura for providing anti-sfCdc25 and anti-sfAkt C-terminus antibodies. We also thank K. Yura for his contribution to perform alignment of SGK isoforms. This work was supported by JSPS KAKENHI grant numbers 17K07405, Takeda Science Foundation, and the Cooperative Program provided by The Atmosphere and Ocean Research Institute, The University of Tokyo.

## Author contributions

E.H., D.H., and K.C. conceived and designed all the experiments, and E.H. performed all the experiments. Cloning of cDNA encoding sfSGK was performed by S.O. and N.H. All authors discussed the results. E.H., D.H., T.K., and K.C. wrote the manuscript.

## Materials and correspondence

Correspondence and requests for materials should be addressed to K.C.

## References

1. Baltz, J. M. Intracellular pH regulation in the early embryo. Bioessays 15, 523–530 (1993).

2. Schatten, G., Bestor, T., Balczon, R., Henson, J. & Schatten, H. Intracellular pH shift initiates microtubule-mediated motility during sea urchin fertilization. Ann. N. Y. Acad. Sci. 466, 940–944 (1986).

3. Grandin, N. & Charbonneau, M. Cycling of intracellular pH during cell division of Xenopus embryos is a cytoplasmic activity depending on protein synthesis and phosphorylation. J. Cell Biol. 111, 523–532 (1990).

4. Grillo-Hill, B., Choi, C., Jimenez-Vidal, M. & Barber, D. Increased H^+^ efflux is sufficient to induce dysplasia and necessary for viability with oncogene expression. Elife 4, 10.7554/eLife.03270. (2015).

5. Orlowski, J. & Grinstein, S. Diversity of the mammalian sodium/proton exchanger SLC9 gene family. Pflugers Archi. 447, 549–565 (2004).

6. He, P. et al. Serum- and glucocorticoid-induced kinase 3 in recycling endosomes mediates acute activation of Na^+^/H^+^ exchanger NHE3 by glucocorticoids. Mol. Biol. Cell 22, 3812–3825 (2011).

7. Wang, D., Sun, H., Lang, F. & Yun, C. Activation of NHE3 by dexamethasone requires phosphorylation of NHE3 at Ser663 by SGK1. Am J Physiol. Cell Physiol. 289, C802–C810 (2005).

8. Yun, C. C. Concerted roles of SGK1 and the Na^+^/H^+^ exchanger regulatory factor 2 (NHERF2) in regulation of NHE3. Cell Physiol. Biochem. 13, 29–40 (2003).

9. Takahashi, E. et al. p90^RSK^ is a serum-stimulated Na^+^/H^+^ exchanger isoform-1 kinase. Regulatory phosphorylation of serine 703 of Na^+^/H^+^ exchanger isoform-1. J. Biol. Chem. 274, 20206–20214 (1999).

10. Wang, C. et al. Somatostatin stimulates intestinal NHE8 expression via p38 MAPK pathway. Am. J. Physiol. Cell Physiol. 300, C375–C382 (2011).

11. Xu, H., Ghishan, F. & Kiela, P. SLC9 gene family: function, expression, and regulation. Copr. Physiol. 8, 555–583 (2018).

12. Dubé, F. & Eckberg, W. Intracellular pH increase driven by an Na^+^/H^+^ exchanger upon activation of surf clam oocytes. Dev. Biol. 190, 41–54 (1997).

13. Lee, S. & Steinhardt, R. pH changes associated with meiotic maturation in oocytes of Xenopus laevis. Dev. Biol. 85, 358–369 (1981).

14. Rezai, K., Kulisz, A. & Wasserman, W. J. Protooncogene product, c-mos kinase, is involved in upregulating Na^+^/H^+^ antiporter in Xenopus oocytes. Am. J. Physiol. 267, C1717–1722 (1994).

15. Miyazaki SI, Ohmori H, & Sasaki S. Potassium rectifications of the starfish oocyte membrane and their changes during oocyte maturation. J. Physiol. 246, 55–78. (1975)

16. Harada, K., Oita, E. & Chiba, K. Metaphase I arrest of starfish oocytes induced via the MAP kinase pathway is released by an increase of intracellular pH. Development 130, 4581–4586 (2003).

17. Moriwaki, K., Nakagawa, T., Nakaya, F., Hirohashi, N. & Chiba, K. Arrest at metaphase of meiosis I in starfish oocytes in the ovary is maintained by high CO_2_ and low O_2_ concentrations in extracellular fluid. Zoolog. Sci. 31, 975–984 (2013).

18. Masui, Y. From oocyte maturation to the *in vitro* cell cycle: the history of discoveries of Maturation-Promoting Factor (MPF) and Cytostatic Factor (CSF). Differentiation 69, 1–17 (2001).

19. Hunt, T. Maturation promoting factor, cyclin and the control of M-phase. Curr. Opin. Cell Biol. 1, 268–274 (1989).

20. Nurse, P. Universal control mechanism regulating onset of M-phase. Nature 344, 503–508 (1990).

21. Kishimoto, T. Entry into mitosis: a solution to the decades-long enigma of MPF. Chromosoma 124, 417–428 (2015).

22. Chiba, K. Evolution of the acquisition of fertilization competence and polyspermy blocks during meiotic maturation. Mol. Reprod. Dev. 78, 808–813 (2011).

23. Kishimoto, T. MPF-based meiotic cell cycle control: Half a century of lessons from starfish oocytes. Proc. Jpn. Acad. Ser. B. Phys. Biol. Sci. 94, 180–203 (2018).

24. Kanatani, H., Shirai, H., Nakanishi, K. & Kurokawa, T. Isolation and indentification of meiosis inducing substance in starfish *Asterias amurensis*. Nature 221, 273–274 (1969).

25. Harada, K., Fukuda, E., Hirohashi, N. & Chiba, K. Regulation of intracellular pH by p90Rsk-dependent activation of an Na^+^/H^+^ exchanger in starfish oocytes. J. Biol. Chem. 285, 24044–24054 (2010).

26. Oita, E., Harada, K. & Chiba, K. Degradation of polyubiquitinated Cyclin B is blocked by the MAPK pathway at the Metaphase I arrest in starfish oocytes. J. Biol. Chem. 279, 18633–18640 (2004).

27. Usui, K., Hirohashi, N. & Chiba, K. Involvement of mitogen-activating protein kinase and intracellular pH in the duration of the metaphase I (MI) pause of starfish oocytes after spawning. Dev. Growth Differ. 50, 357–364 (2008).

28. Ochi, H., Aoto, S., Tachibana, K., Hara, M. & Chiba, K. Block of CDK1-dependent polyadenosine elongation of Cyclin B mRNA in metaphase-I-arrested starfish oocytes is released by intracellular pH elevation upon spawning. Mol. Reprod. Dev. 83, 79–87 (2016).

29. Shilling, F., Chiba, K., Hoshi, M., Kishimoto, T. & Jaffe, L. Pertussis toxin inhibits 1-methyladenine-induced maturation in starfish oocytes. Dev. Biol. 133, 605–608 (1989).

30. Chiba, K., Kontani, K., Tadenuma, H., Katada, T. & Hoshi, M. Induction of starfish oocyte maturation by the bg subunit of starfish G protein and possible existence of the subsequent effector in cytoplasm. Mol. Biol. Cell 4, 1027–1034 (1993).

31. Sadler, K. & Ruderman, J. Components of the signaling pathway linking the 1-methyladenine receptor to MPF activation and maturation in starfish oocytes. Dev. Biol. 197, 25–38 (1998).

32. Hiraoka, D., Aono, R., Hanada, S., Okumura, E. & Kishimoto, T. Two new competing pathways establish the threshold for cyclin-B–Cdk1 activation at the meiotic G2/M transition. J. Cell Sci. 129, 3153–3166 (2016).

33. Vanhaesebroeck, B., Guillermet-Guibert, J., Graupera, M. & Bilanges, B. The emerging mechanisms of isoform-specific PI3K signalling. Nat. Rev. Mol. Cell Biol. 11, 329–341 (2010).

34. Tessier, M. & Woodgett, J. Serum and glucocorticoid-regulated protein kinases: Variations on a theme. J. Cell Biochem. 98, 1391–1407 (2006).

35. Kobayashi, T., Deak, M., Morrice, N. & Cohen, P. Characterization of the structure and regulation of two novel isoforms of serum- and glucocorticoid-induced protein kinase. Biochem. J. 344, 189–197 (1999).

36. Malik, N. et al. Mechanism of activation of SGK3 by growth factors via the Class 1 and Class 3 PI3Ks. Biochem. J. 475, 117–135 (2018).

37. Pearce, L., Komander, D. & Alessi, D. The nuts and bolts of AGC protein kinases. Nat. Rev. Mol. Cell Biol. 11, 9–22 (2010).

38. Kobayashi, T. & Cohen, P. Activation of serum- and glucocorticoid-regulated protein kinase by agonists that activate phosphatidylinositide 3-kinase is mediated by 3-phosphoinositide-dependent protein kinase-1 (PDK1) and PDK2. Biochem. J. 339, 319–328 (1999).

39. Tessier, M. & Woodgett, J. R. Role of the Phox homology domain and phosphorylation in activation of serum and glucocorticoid-regulated kinase-3. J. Biol. Chem. 281, 23978–23989 (2006).

40. García-Martínez, J. M. & Alessi, D. R. mTOR complex 2 (mTORC2) controls hydrophobic motif phosphorylation and activation of serum- and glucocorticoid-induced protein kinase 1 (SGK1). Biochem. J. 416, 375–385 (2008).

41. Biondi, RM., Kieloch, A., Currie, RA., Deak, M. & Alessi, DR. The PIF-binding pocket in PDK1 is essential for activation of S6K and SGK, but not PKB. EMBO J. 20, 4380–4390 (2001).

42. Lu, M. et al. mTOR Complex-2 Activates ENaC by Phosphorylating SGK1. J. Am. Soc. Nephrol. 21, 811–818 (2010).

43. Lien, E. C., Dibble, C. C. & Toker, A. PI3K signaling in cancer: beyond AKT. Curr. Opin. Cell Biol. 45, 62–71 (2017).

44. Hiraoka, D. et al. PDK1 is required for the hormonal signaling pathway leading to meiotic resumption in starfish oocytes. Dev. Biol. 276, 330–336 (2004).

45. Hiraoka, Okumura & Kishimoto, T. Turn motif phosphorylation negatively regulates activation loop phosphorylation in Akt. Oncogene 30, 4487–4497 (2011).

46. Okumura, E. et al. Akt inhibits Myt1 in the signalling pathway that leads to meiotic G2/M-phase transition. Nat. Cell. Biol. 4, 111–116 (2002).

47. Okumura, E., Sekiai, T., Hisanaga, S., Tachibana, K. & Kishimoto, T. Initial triggering of M-phase in starfish oocytes: a possible novel component of maturation-promoting factor besides cdc2 kinase. J. Cell Biol. 132, 125–135 (1996).

48. Eltschinger, S. & Loewith, R. TOR complexes and the maintenance of cellular homeostasis. Trends Cell Biol. 26, 148–159 (2016).

49. Kinoshita, E., Kinoshita-Kikuta, E., Takiyama, K. & Koike, T. Phosphate-binding Tag, a New Tool to Visualize Phosphorylated Proteins. Mol. Cell Proteomics 5, 749–757 (2006).

50. Miyazaki, S. & Hirai, S. Fast polyspermy block and activation potential correlated changes during oocyte maturation of a starfish. Dev. Biol. 70, 327–340 (1979).

51. Hayashi, M. et al. BMK1 mediates growth factor-induced cell proliferation through direct cellular activation of SGK. J. Biol. Chem. 276, 8631–8634 (2001).

52. Gasser, J. et al. SGK3 mediates INPP4B-dependent PI3K signaling in breast cancer. Mol. Cell 56, 595–607 (2014).

53. Hiraoka, D., Hosoda, E., Chiba, K. & Kishimoto, T. SGK phosphorylates Cdc25 and Myt1 to trigger cyclin B-Cdk1 activation at the meiotic G2/M transition. Preprint at https://doi.org/10.1101/499640 (2018).

54. Terasaki, M. et al. A new model for nuclear envelope breakdown. Mol. Biol. Cell 12, 503–510 (2001).

55. Lénárt, P. et al. Nuclear envelope breakdown in starfish oocytes proceeds by partial NPC disassembly followed by a rapidly spreading fenestration of nuclear membranes. J. Cell Biol. 160, 1055–1068 (2003).

56. Mori, M. et al. An Arp2/3 nucleated F-actin shell fragments nuclear membranes at nuclear envelope breakdown in starfish oocytes. Curr. Bio. 24, 1421–1428 (2014).

57. Jenkins, E., Debnath, S., Varriano, S., Gundry, S. & Fata, J. Na^+^/H^+^ exchanger 1 (NHE1) function is necessary for maintaining mammary tissue architecture. Dev. Dyn. 243, 229–242 (2014).

58. Rodeau, J. L. & Vilain, J. P. Changes in membrane potential, membrane resistance, and intracellular H^+^, K^+^, Na^+^, and Cl^−^ activities during the progesterone-induced maturation of urodele amphibian oocytes. Dev. Biol. 120, 481–493 (1987).

59. Stith, B. & Maller, J. Increased intracellular pH is not necessary for ribosomal protein S6 phosphorylation, increased protein synthesis, or germinal vesicle breakdown in Xenopus oocytes. Dev. Biol. 107, 460–469 (1985).

60. Sellier, C. et al. Intracellular acidification delays hormonal G2/M transition and inhibits G2/M transition triggered by thiophosphorylated MAPK in Xenopus oocytes. J. Cell Biochem. 98, 287–300 (2006).

61. Tachibana, K., Machida, T., Nomura, Y. & Kishimoto, T. MAP kinase links the fertilization signal transduction pathway to the G1/S-phase transition in starfish eggs. EMBO J. 16, 4333–4339 (1997).

62. Tachibana, K., Tanaka, D., Isobe, T. & Kishimoto, T. c-Mos forces the mitotic cell cycle to undergo meiosis II to produce haploid gametes. Proc. Natl. Acad. Sci. USA 97, 14301–14306 (2000).

63. Flament, S., Browaeys, E., Rodeau, J. L., Bertout, M. & Vilain, J. P. Xenopus oocyte maturation: cytoplasm alkalization is involved in germinal vesicle migration. Int. J. Dev. Biol. 40, 471–476 (1996).

64. Kishimoto, T. Microinjection and cytoplasmic transfer in starfish oocytes. Methods Cell Biol. 27, 379–394 (1986).

65. Chiba, K. et al. The primary structure of the α subunit of a starfish guanosine-nucleotide-binding regulatory protein involved in 1-methyladenine-induced oocyte maturation. Eur. J. Biochem. 207, 833–838 (1992).

66. Hamaguchi, M., Watanabe, K. & Hamaguchi, Y. Regulation of intracellular pH in sea urchin eggs by medium containing both weak acid and base. Cell Struct. Funct. 22, 387–398 (1997).

67. Hijikata, A., Yura, K., Noguti, T. & Go, M. Revisiting gap locations in amino acid sequence alignments and a proposal for a method to improve them by introducing solvent accessibility. Proteins 79, 1868–1877 (2011).

