## Supplementary figures for "SGK regulates pH increase and cyclin B-Cdk1 activation to resume meiosis in starfish ovarian oocytes"

**Supplementary Information**

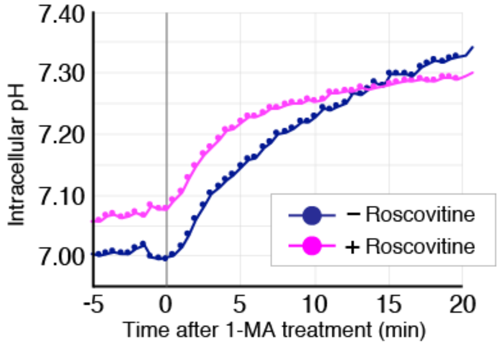


**Supplementary Figure 1  pH$_i$ increase soon after 1-MA stimulus is independent of cyclin B–Cdk1 activation.**

Unstimulated oocytes were injected with BCECF-dextran, incubated in the presence or absence of the selective Cdk1 inhibitor roscovitine for 1 h, and then, treated with 1-MA. Fluorescence intensity ratio was measured every 30 seconds before and after 1-MA addition. Then, pH$_i$ was calculated based on the fluorescence intensity ratio. Average pH$_i$ values of 11 and 8 oocytes in the presence and absence of roscovitine, respectively, are plotted on the indicated graph. This result is a representative of two independent experiments. pH$_i$ was elevated after 1-MA stimulus even in the presence of roscovitine, as in control oocytes.

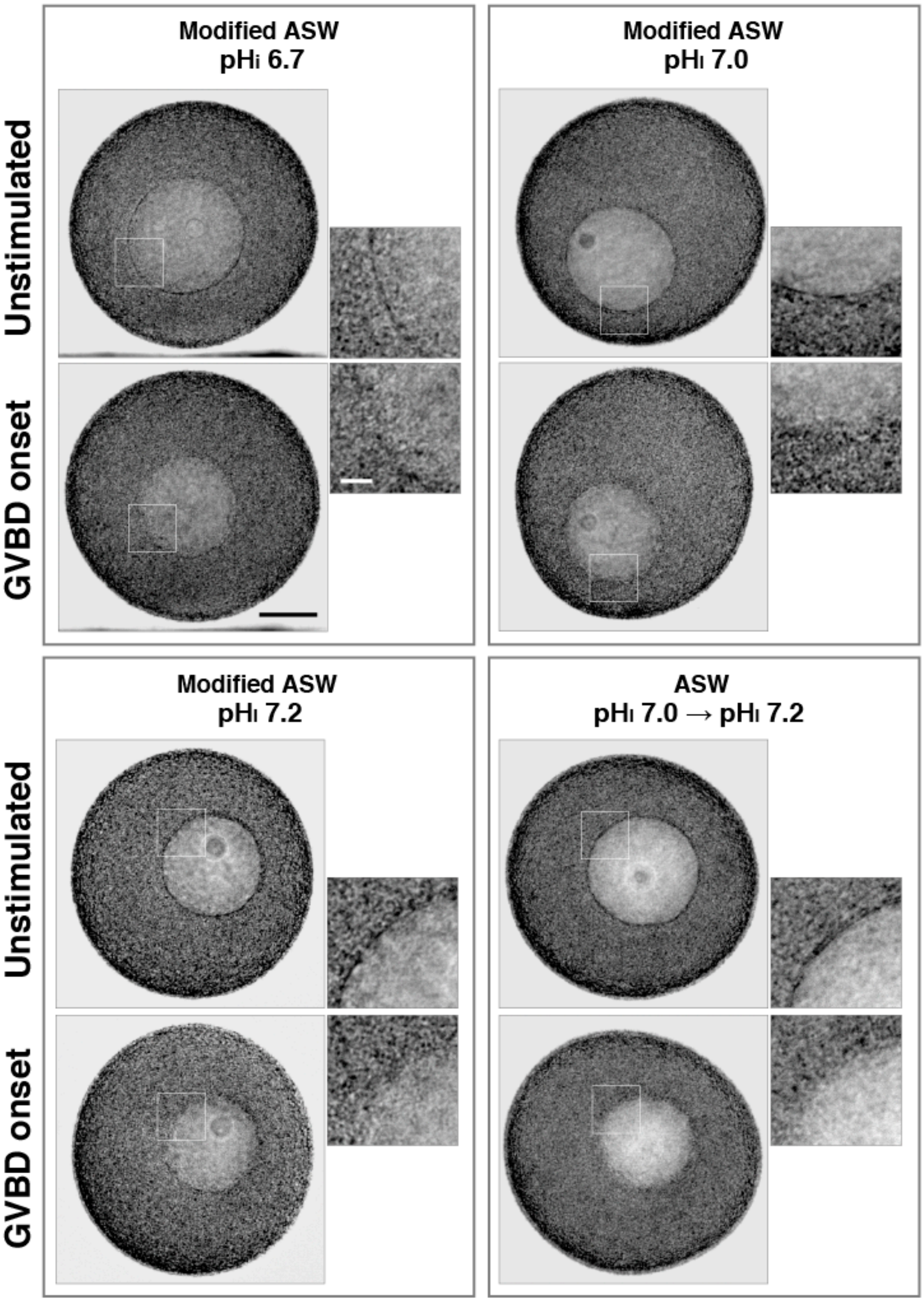


**Supplementary Figure 2 Morphology of GVBD onset is normal at various pHi values.**

To clamp $pH_i$ at 6.7, 7.0 and 7.2, unstimulated oocytes were incubated with sodium-free artificial seawater containing $CH_3COONH_4$ (Modified ASW) for 20 min. As a control, oocytes were incubated in artificial seawater (ASW) for 20 min, in which $pH_i$ is approximately 7.0, and increase to ~7.2 after 1-MA treatment. Nomarski differential interference contrast (DIC) images of unstimulated oocytes and 1-MA stimulated oocytes at onset of GVBD were captured. Left column: images of whole oocytes. Right column: enlarged view of the rim of GV corresponding to the white rectangles. The black scale bar in the left column represents 50 μm, and the white scale bar in the right column represents 10 μm. Results in all panels are representative of two independent experiments. In ASW, rim of GV in unstimulated oocytes looked clear line. Cytoplasmic granules started to invade into the inner GV area at GVBD onset. Rim of GV in unstimulated oocytes and at GVBD onset at all clamped $pH_i$ values were morphologically indistinguishable from those in ASW.

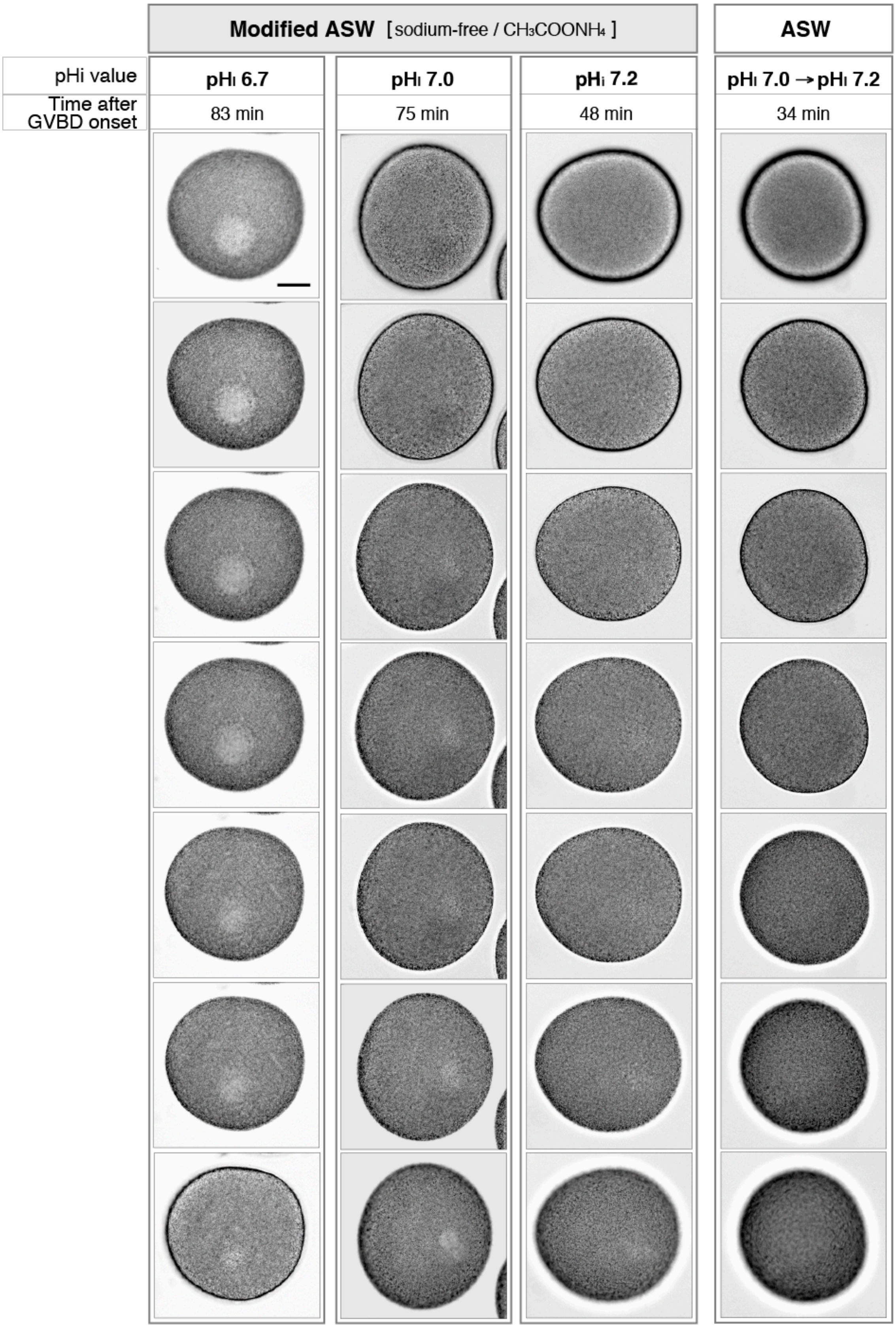


**Supplementary Figure 3 Completion of GVBD was ascertained by Z-stack images.**

Unstimulated oocytes were incubated in modified ASW to clamp $pH_i$ at 6.7, 7.0, and 7.2 or in ASW as a control, and then treated with 1-MA. Time-lapse DIC imaging was performed (see Supplementary Movies 1, 2, 3, and 4). Then, to ascertain whether GVBD was completed, Z-stack images of these oocytes were taken by DIC microscopy at the indicated times after GVBD onset. The scale bar represents 50 µm. Cytoplasmic granules were homogenously distributed throughout oocytes in ASW and oocytes at clamped $pH_i$ of 7.2. At a clamped $pH_i$ of 7.0, a trace of the inner nuclear region was observed. At a $pH_i$ of 6.7, a large part of the inner nuclear region had not been invaded by the granules.

**Supplementary Movies 1–4**

Unstimulated oocytes were incubated in modified ASW to clamp $pH_i$ at 6.7, 7.0, and 7.2, or incubated in ASW as a control. Time-lapse DIC imaging was performed every 10 seconds before and after 1-MA treatment. The movies start at 3 min before 1-MA treatment. “h: m: s” in frames represents hour, minutes and seconds after 1-MA treatment. All images were acquired by focusing the microscope on the equatorial plane of GV region. When the equatorial plane was moved out of focus during the time-lapse imaging, the focus was returned on the equatorial plane. Selected images from the image sequence are shown in Fig. **5c**.

**Movie 1** GVBD of oocytes in ASW

Focus was not changed during time-lapse imaging. This movie runs for 45 min.

**Movie 2** GVBD of oocytes in modified ASW for clamping pHi at 6.7

To focus on the rim of the GV, vertical focus was changed at 0:04:30, 0:20:40, 0:24:00, 0:27:40, 0:33:20, 0:42:00, 0:46:10, 0:57:50, 01:02:50, and 01:07:30. This movie runs for 1 h 20 min.

**Movie 3** GVBD of oocytes in modified ASW for clamping pHi at 7.0

Focus was not changed during time-lapse imaging. This movie runs for 1 h 20 min.

**Movie 4** GVBD of oocytes in modified ASW for clamping pHi at 7.2

Focus was not changed during time- lapse imaging. This movie runs for 55 min.
